# Recognition memory via repetition suppression in mouse hippocampal dorsal CA2 pyramidal neurons expressing the vasopressin 1b receptor

**DOI:** 10.1101/2020.05.11.078915

**Authors:** Adi Cymerblit-Sabba, Michelle Stackmann, Sarah K. Williams Avram, Michael C. Granovetter, Nicholas I. Cliz, Francisco Pereira, Adam S. Smith, June Song, Heon-Jin Lee, W. Scott Young

## Abstract

Recognition memory, often compromised in psychiatric disorders, is a major component of declarative memory, which permits the realization that an event, object or social subject has been previously encountered. The CA2 region of the dorsal hippocampus (dCA2) is involved in social memory and responds to novel objects, in time and space. However, it remains unclear how these neurons encode either social or inanimate object recognition. Here, we show that in dCA2, encoding of social recognition memory entails suppression of pyramidal neurons’ activity leading to a sparse representation of the familiar conspecific. We discuss the neural coding scheme by which dCA2 pyramidal neurons contribute to social memory.

## Introduction

The ability to remember conspecifics is critical for the establishment of social relationships in species living in complex social systems. Rodents show a preference for investigating a novel conspecific over a familiar one^1, 2^. In the lab, social memory can be assessed reliably by measuring the reduction in investigation of a familiar relative to a novel conspecific.

We and others have shown in rodent models that the hippocampal CA2 subfield plays an important role in the acquisition of social memories ^3-6^ and, more specifically, the key involvement of the vasopressin 1b^4, 7-9^ and oxytocin receptors^10, 11^ (Avpr1b and Oxtr, respectively). The findings of the unique characteristics and functionality of the CA2 hippocampal subfield^12, 13^ established it as an intra-hippocampal hub that is also connected with other brain regions - conceivably a key player in the integration of social, spatial and temporal inputs, with possibly distinct subpopulations underlying this ability. Yet, the way CA2 pyramidal neurons encode social inputs remains unclear.

In humans, abnormalities of CA2 neurons have been linked to several psychiatric illnesses, such as schizophrenia, depression and Alzheimer’s disease^14-16^. Moreover, there is mounting evidence of impaired declarative memory processes in patients with bipolar disorder^17^ as well as positive correlation between hippocampal volume and severity of psychosis, declarative memory and overall cognitive performance, with most prominent reduction in CA2/3^18^. Thus, deciphering the neuronal substrates of social memory is of great importance.

In this study, we used *in vivo* calcium imaging to visualize the dynamic of dorsal CA2 (dCA2) pyramidal neurons (the vast majority, if not all, express Avpr1b as described below) underlying social memory in Avpr1b^+/Cre^ transgenic mice. We demonstrate how the neurons’ activity recapitulates the sniffing behavior exhibited by the imaged mice toward a familiar conspecific, as well as the creation of sparse neuronal representation of it, which is absent for the inanimate object. These results suggest the existence of parallel processing with enhanced salience of social cues in dCA2 pyramidal neurons.

## Materials and Methods

Materials and Methods are available in the Supplemental Information.

## Results

### Validation of transgenic mouse model

We generated a knock-in (KI) mouse line in which Cre expression is driven by the Avpr1b promoter (detailed previously^22^). At the same level as our lens implant at rostral dCA2, *in situ* hybridization histochemistry demonstrates the expression of Avpr1b in the CA2 as well as in the immediately adjacent and interdigitating border with the CA3 region where the marker Bok is distinctly expressed (Supplemental Fig. 1a-b,e; Supplemental Table 1). As we described previously^22^, Avpr1b expression remains confined to the CA2 subfield along the dorsal-ventral axis (more ventral in Supplemental Fig. 1c,d). Moreover, the expression patterns are identical to those seen with the accepted CA2 markers Pcp4 (Supplemental Fig.1d-e), Map3k15 (Supplemental Fig.1f-g) and Amigo 2 (Supplemental Fig.1 j-l) that, of course, also interdigitate with CA3. Consistent with these current observations and our previous Avpr1b localization^6^, Cre recombinase expression is also confined to the CA2 and immediately adjacent interdigitating CA3 regions (Supplemental Fig. 2a) where it is expressed in 87.3% of the neurons in the pyramidal cell layer (over 9% are interneurons ^23^) (Supplemental Fig. 2b-e). This permits delivery of the virally expressing Cre-activatable GCaMP6s specifically into the dCA2 pyramidal neurons for subsequent imaging that is restricted to dCA2. Behaviorally, the heterozygotes (Avpr1b^+/Cre^) mice show no deficit in social memory^22^ while homozygous KI (Avpr1b^Cre/Cre^) show a deficit in social recognition (detailed description in Supplemental Fig. 3) similarl to our first Avpr1b knock out^9^.

### Sparse neuronal representation in dCA2 pyramidal neurons

Previous data, including differential gene expression and connectivity patterns^24^, support the different roles of neuronal populations along the dorsoventral axis of the hippocampus in mnemonic and emotional functions^25, 26^. Here, we used cellular-resolution calcium imaging to visualize the activity of rostral dCA2 neurons in Avpr1b^+/Cre^ mice during memory acquisition and short-term retrieval. A gradient-refractive lens and a head-mounted mini-microscope (Fig. 1a-b) allowed observation of the calcium dynamics while our mice were performing a battery of behavioral paradigms involving recognition memory and novelty detection (Supplemental Fig. 4), followed by neural data analyses including cell identification, extraction of calcium traces, calcium event detection and comparison of neuronal activity across trials (Fig. 1c-e).

**Figure 1.**
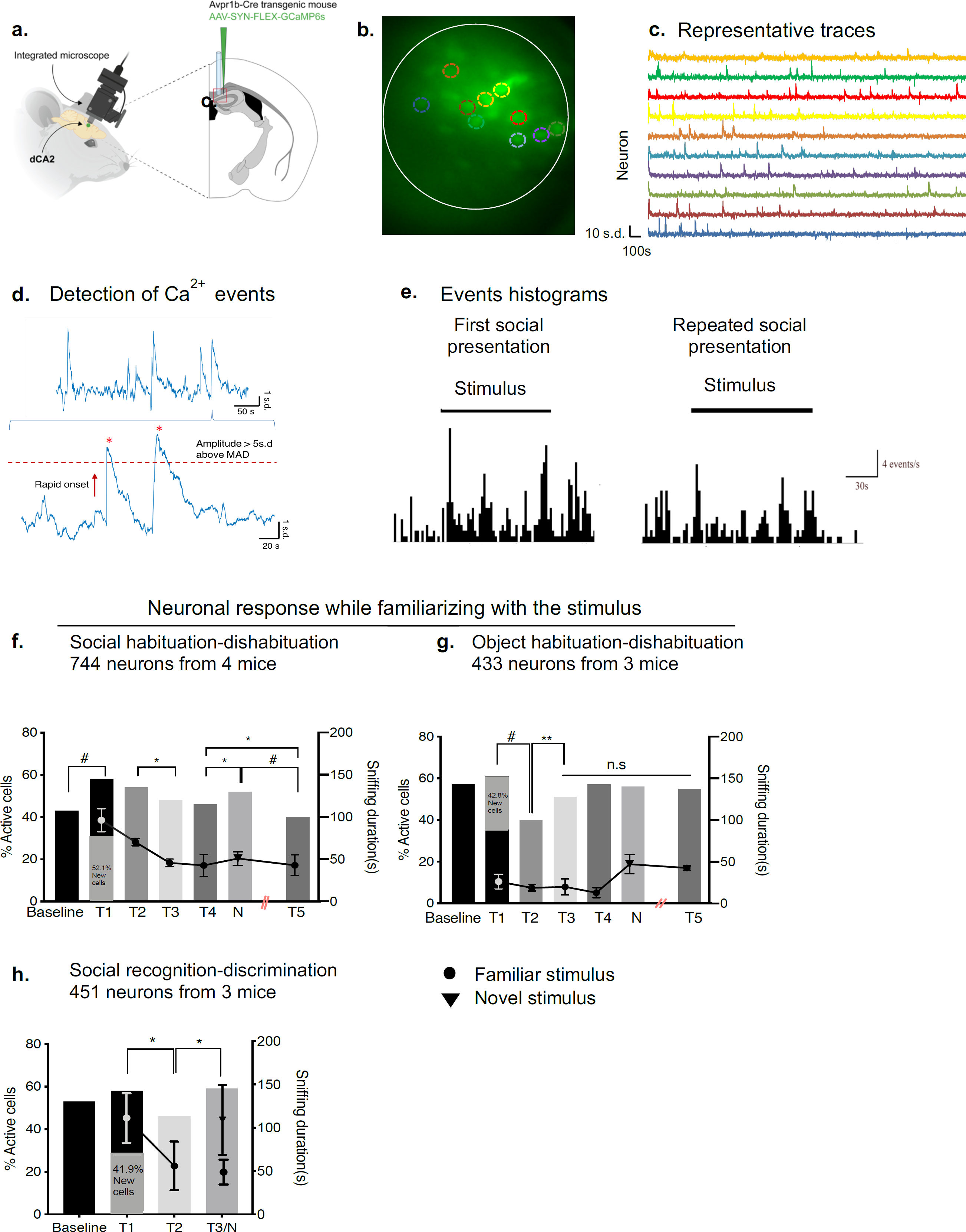
Visualizing pyramidal neurons of dorsal CA2. **a.** Viral delivery of AAV1-Syn-Flex-GCamp6s into dCA2 of Avpr1b+/Cre mice. **b.** representative field of view. **c**. Representative traces recorded from 10 dCA2 pyramidal neurons during the performance of the social habituation-dishabituation (SHD) paradigm. **d**. Calcium events were detected based on defined kinetic criteria, with a 5-standard deviation increase above the median absolute deviation of the trace. **e.** Representative histograms of calcium events across the first and fourth social exposures in SHD paradigm. **f.** The percentage of active neurons across presentations of the social stimulus (bar graphs, Fisher’s exact test: Baseline-T1 *P*<0.0001, T2-T3 *P*<0.03, T4-N *P*<0.03, N- T5 *P*<0.0001, T4-T5, *P*<0.03), and the sniffing behavior of the imaged mice (line graph).a. **g**. The percentage of active neurons across presentations of the object stimulus (bar graphs, Fisher’s exact test: Baseline-T1 *P*>0.2, T1-T2 *P*<0.0001; T2-T3 *P*<0.002), and the sniffing behavior of the imaged mice (line graph). **h**. The percentage of active neurons across recognition and discrimination of social stimuli (bar graphs, Fisher’s exact test: Baseline-T1 *P*<0.2, T1-T2 *P*<0.0005, T2-T3/N *P*<0.0001), and the sniffing behavior of the imaged mice (line graph). Data as mean±sem.

The sparse coding view argues that a small neuronal ensemble responds in an explicit manner to specific features, objects or concepts^27^, and carries a large amount of information due to previous computations. Moreover, sparsification and selective neuronal activation have been described in the human temporal lobe upon familiarization with a stimulus^27, 28^. To determine whether dCA2 pyramidal neurons employ a coding scheme for familiarized stimuli that involves selective neuronal activation, we compared the number of active neurons across trials of the behavioral paradigms.

From the 744 neurons collected for the social habituation-dishabituation (SHD) paradigm, a great percentage of neurons was active in the first presentation of the conspecific (T1-stim 1)(Fig. 1f). Moreover, 52.1% of them were silent at baseline (new cells). The percentage of active neurons decreased along the repeating trials of the familiar conspecific (T2-T4; stim 1), then rose upon the presentation of second presentation of a novel conspecific (N-stim 2). In the final presentation of the familiar conspecific (T5- stim 1), following a longer interval of ^45-60^ min, a decreased percentage of responding neurons was observed compared to the previous familiar or novel presentation. The percentage of the active cells across trials recapitulated the sniffing behavior of the imaged mice toward the conspecific presented to them (Fig. 1f, lined graph).

From the 433 neurons collected for the object habituation-dishabituation (OHD) paradigm, we did not observe an increase in the percentage of active neurons in the first presentation of the novel object (Fig. 1g). Still, over 40% of the active neurons in the first trial were silent at baseline (new cells). Except for a drop in the neurons’ response in the second trial (T2), the percentage of active neurons remained stable across the following presentations of either the familiar or the second novel object. The low sniffing durations shown by the imaged mice in the first presentation of the object (Fig. 1g, line graph) did not result in lower neuronal response, but actually a similar percentage of active neurons as in the presence of the social stimulus. Moreover, in the fourth trial of the object paradigm, while the mice decreased their sniffing duration, the number of active neurons was similar to the that seen in the first and novel trials, in which they spent more time sniffing the object.

Finally, from the 451 neurons collected for the social recognition-discrimination (SRD) paradigm, the percent of active neurons in the first presentation of the novel conspecific did not increase (Fig. 1h), but over 40% of the neurons were silent at baseline (new cells). Following a 30-min interval, a decreased percentage of active neurons was observed in the recognition trial. Following a second 30-min interval, in the discrimination trial, with both the familiar and novel stimuli in the cage, more neurons were active, again recapitulating the sniffing behavior toward the conspecifics presented.

While the changes seem to be stimulus dependent, we were not able to identify selective neurons responding to a specific conspecific.

The percentages of active neurons across the inter-trial intervals in all three paradigms, when the stimuli were removed from the cage, were different (Supplemental Fig. 5a-c, left graphs; %Active cells). In the social habituation-dishabituation paradigm, a lower percentage of active neurons was observed in the last two intervals and no changes were observed in the SRD. A lower percent of active neurons was observed in the second and fourth intervals of OHD. Taken together, the changes in the number of active neurons across the trials appear to be stimulus-dependent consistent with neuronal sparsification during social recognition and no change in the representation of the object.

### dCA2 pyramidal neurons are more active in the presence of a social stimulus

In rats, spatial representation in CA2 neurons was shown to remap following an exposure to either novel social or object stimuli, with no effect on the averaged firing rate^5^. In mice, GCaMP6f signal in dCA2 increased during exploration of both a novel female or male^29^. Moreover, the population activity tended to decrease along familiarization with the conspecific, with no changes in a novel environment or in the presence of novel object. The inconsistency could arise from methodological differences. The cellular-resolution visualization of rostral dCA2 neurons in our Avpr1b^+/Cre^ mice allowed us to extend the understanding of how this region encodes social and non-social familiarization.

For every neuron we identified in the social and object habituation-dishabituation paradigms, we compared the total number of calcium events generated by it in the 2-min trial, in the presence or absence of the stimulus (Fig. 2 a-b, scatterplots). The majority of the neurons increased their calcium activity upon the first presentation of the novel conspecific (Fig. 2a- first row). Moreover, the majority of the neurons generated more calcium events in the presence of the familiar conspecific compared with times when it was removed from the cage (Fig. 2a- second row;T5, third row) as well as in the presence of the second novel conspecific (N, Fig. 2a- third row). While the majority of the neurons increased their activity upon the first presentation of the novel object (Fig. 2b- first row), there was no change when the object was removed from the cage in the following inter-trial interval. Also, there were no changes in neuronal responses between trials whether the object was present or not in the cage (T2,T3, Fig. 2b- second row) except in the fourth presentation (T4) when a subset of the neurons decreased their activity when the object was removed from the cage (Fig. 2b- second row). We did not observe a significant increase in neuronal activity when mice were presented again with a novel object (N, Fig. 2b- third row) nor in the final presentation of the familiar object (Fig. 2b- third row). Yet, the scatterplots suggest that a subset of neurons do respond to the presence of the object in the cage.

**Figure 2.**
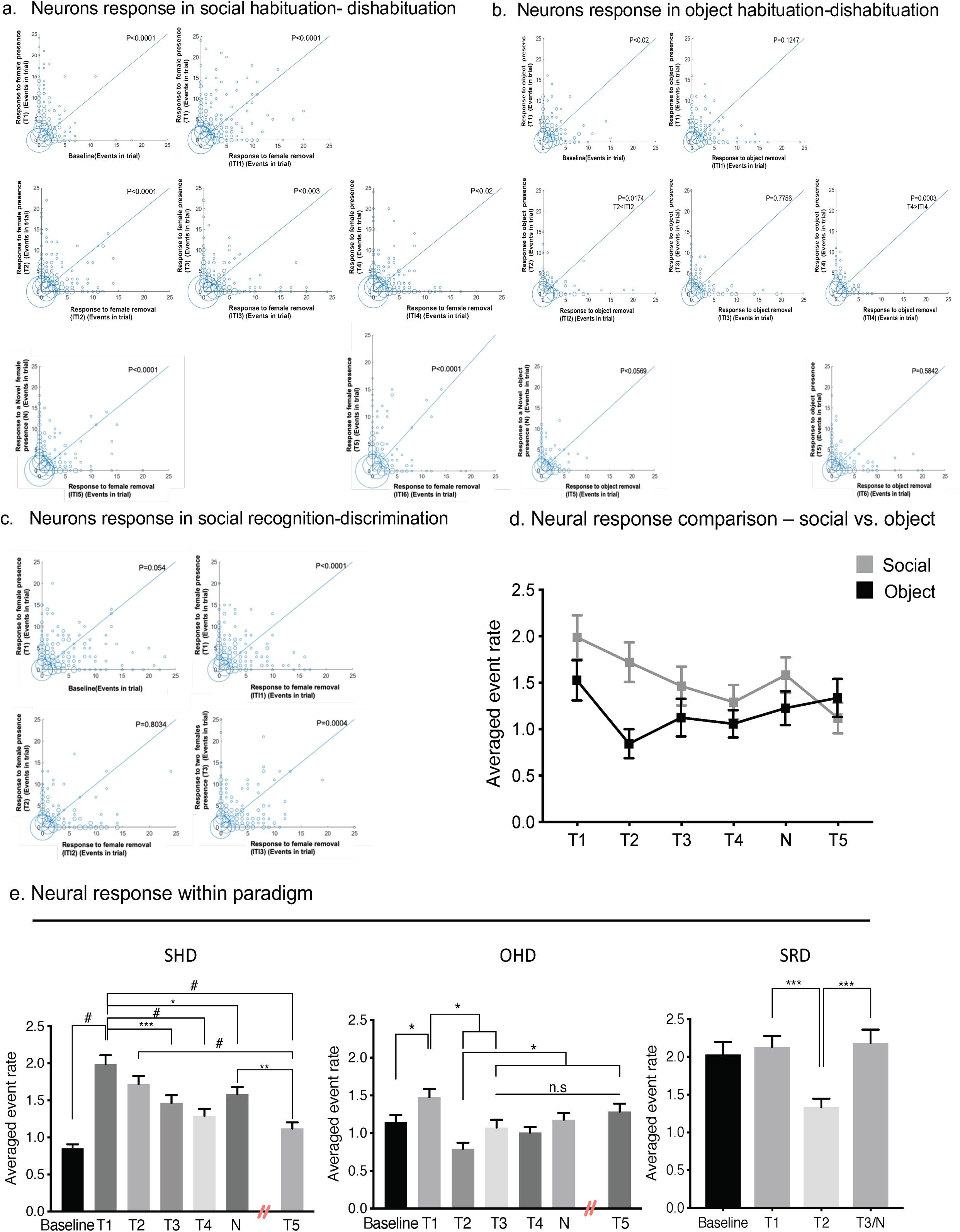
Neuronal response to presence of a stimulus. **a.** Scatter plot comparisons of neuronal responses to the presence and removal of the social stimulus, across all trials of the SHD. Neurons had a greater response to the presence of the social stimulus in the cage. **b.** Scatter plots comparisons of neuronal responses to the presence and removal of the object stimulus, across all trials of OHD. Neurons responded to the first presentation of the object (T1 vs. Baseline). A decreased response was observed only following the fourth presentation of the object. **c**. Scatter plots comparisons of neuronal responses to the presence and removal of the social stimulus, across all trials of the SRD. Neurons decreased their reponses upon removal of the social stimulus in the first and last trials (T1 vs. ITI1 and T3 vs. ITI3). The size of the bubbles represent the number of neurons with similar calcium events in the compared trials. Wilcoxon matched-pairs signed rank test with Pratt method was used to compare trials. **d.** Neurons’ calcium event rates in response to the social versus the object stimulus presentations (SHD: n=744, OHD: n=433 from 5 mice: generalized linear mixed model: stimulus X neurons, *P*<0.0001). **e.** (Left bars) The calcium event rate gradually decreased across social repetitions (SHD: n=744 neurons from 4 mice: Wilcoxon signed rank test Baseline-T1 *P*<0.0001; repeated measures Friedman test: neurons X trials *P*<0.0001 with Dunn’s multiple comparisons test T1-T3 *P*<0.03, T1-T4 *P*<0.0003, T1-T5 *P*<0.0001, T2- T5 *P*<0.002, T3-T5 *P*<0.02, N-T5 *P*<0.0002).(Middle bars) Calcium event rates across the object repetitions remained consistent following the second trial (OHD: n=433 neurons from 3 mice; Wilcoxon signed rank test for Baseline-T1, *P*<0.02; repeated measures Friedman test: neurons X trials, *P* < 0.0001; Dunn’s multiple comparisons test: T2:T1,T4,N,T5 *P*<0.02). (Right bars) Suppression of calcium event rates was observed in the second trial followed by increased rate upon novelty presentation (SRD: n=451 neurons from 3 mice; Wilcoxon signed rank test Baseline-T1 *P*>0.08; repeated measures Friedman test: neurons X trials *P*<0.0001; Dunn’s multiple comparison: T1-T2 *P*<0.0001; T2-T3/N *P*<0.0005). Bars shown as mean ± sem: \**,P* <0.05; **,*P* <0.001; ***,*P* <0.0002; #,*P* <0.0001. In all tests, the neuronal data was not distributed normally (D’Agostino & Pearson test P<0.0001). Averaged event rate reflects calcium events in the 2-min trial across all imaged neurons.

Our third behavioral task aimed to examine the changes in the neuronal response to the social stimulus with extended intervals as well a more challenging social task in which they are required to actively discriminate between the familiar and novel conspecifics. The baseline averaged calcium activity in the 451 neurons collected was unexpectedly high even though all three baseline situations were identical. While there was no significant increase in neurons’ activity in the first presentation of the conspecific (Fig. 2c, first row), we did find that over 40% of the active neurons in this trial were silent at baseline. This suggests that the neurons do respond to the presence of the novel conspecific. Moreover, upon its removal, decreased activity was observed in the majority of the neurons (Fig. 2c, first row). The lack of change in neuronal activity in the comparison of the second presentation of the familiar conspecific (T2) to the following interval when it is removed is likely the result of already decreased T2 response upon familiarization. The distribution of the scatter plot suggests the existence of a subset of neurons responsive selectively to the conspecific. Importantly, in the third trial (T3/N), when imaged mice were presented with both the familiar and a novel conspecific, the majority of the neurons show increased activity (Fig. 2c, second row).

An individual animal-based analysis approach allowed us to determine whether the pattern of neuronal activity was reliably different between trials with the stimulus and inter-trial intervals (in the absence of stimulus). In all but one animal, we were able to distinguish between the different trials with significant accuracy (Table 1).

**Table 1.** Summarized accuracy results and active bins included for distinguishing pattern of neuronal activity between trials with the stimulus and inter-trial intervals (ITI, stimulus is absence) and significance at the corrected p=0.01 level. The animal number refers to the individual mouse.

| Test<br>(animal #) | Accuracy | Significance<br>($P<0.01$) | # Bins with<br>Activity |
| --- | --- | --- | --- |
| OHD (1) | 0.66 | Yes | 778 |
| OHD (5) | 0.56 | No | 1312 |
| OHD (4) | 0.66 | Yes | 1152 |
| SHD (2) | 0.67 | Yes | 1068 |
| SHD (3) | 0.69 | Yes | 1748 |
| SHD (1) | 0.65 | Yes | 842 |
| SHD (4) | 0.61 | Yes | 1226 |
| SRD (3) | 0.73 | Yes | 978 |
| SRD (1) | 0.67 | Yes | 458 |
| SRD (4) | 0.70 | Yes | 234 |

In all trials, neuronal activity was not normally distributed. We used a generalized linear mixed model to compare the changes in neurons’ calcium events across the trials’ exposures between the social and object habituation-dishabituation paradigms. We found a significant difference in the pattern of response to the different stimuli (Fig. 2d) as well as increased response in the presence of the social stimulus.

### Repetition suppression in dCA2 pyramidal neurons

In rodents, recognition of a repeatedly presented stimulus results in decreased interest which is indicated by reduced time spent sniffing or exploring the stimulus. While CA2 pyramidal neurons have been shown to code place fields, consistent activity patterns were reported between consecutive sessions in the shaped box^30^. Moreover, in this study, firing rate variability was not detected over short time intervals^30^. One of the major goals in memory research is to understand how it is represented in transient dynamic patterns at the neuronal population level. In order to investigate how patterns of dCA2 neuronal activity change as a novel stimulus becomes familiar in the same cage, we compared the calcium event rate for all collected neurons across the repeated exposures to the stimulus, including silent ones. While an increased rate was observed upon the first presentation of both the social and the object stimuli (Fig. 2e), in the SHD test the calcium event rate gradually declined with familiarization and slightly increased when a novel social stimulus (N) was presented (Fig. 2e, left bar graph). In the OHD test, we observed a decreased calcium event rate in the second exposure, followed by a steady rate across the subsequent exposures to either the familiar (T3-T5) or novel (N) objects (Fig. 2e, middle). In the extended intervals paradigm, SRD, we observed a decreased rate in the recognition trial and a great increase in the subsequent discrimination trial (T3/N) (Fig. 2e, right bar graph). In order to determine whether sniffing information exists in the neuronal activity, we used a machine classifier analysis. This analysis demonstrated that the neuronal activity does not contain information about sniffing (Table 2).

**Table 2.**
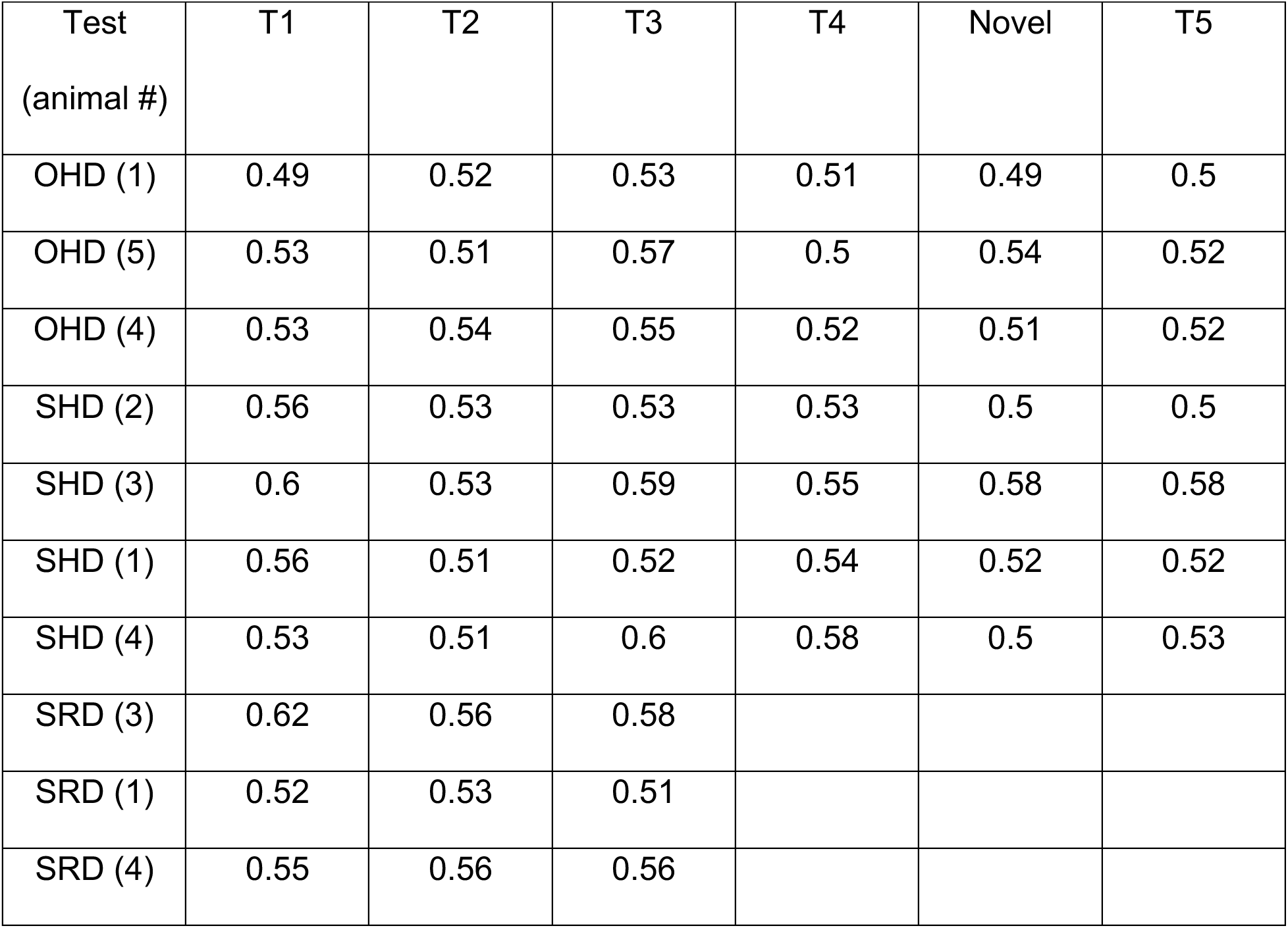
Summarized accuracy results of sniffing by neuronal activity. Accuracy tested using a binomial test with a null hypothesis of classifier chance performance (accuracy=0.5). Sniffing could not be predicted by the neuronal activity. The animal number refers to the individual mouse.

A confound to our observations could arise from the mice changing their mobility during the trials. A subset of pyramidal neurons in CA2 were shown to increase their activity during immobility^31^. Our imaged mice decreased their immobility duration in the first exposure and showed consistent durations of immobility during the following trials (Supplementary Fig. 4f). Thus, we argue the changes we observed were not driven by this unique subset of cells.

Moreover, a different pattern of response appeared across the intervals of all paradigms (Supplemental Fig. 5a-c middle graphs). In the SHD test, the rate remained stable until reduced in the last two intervals (ITI5, ITI6). A reduced rate was observed in the fourth interval of the OHD test (ITI4) and no differences were seen in the SRD test. Furthermore, as we described in the scatter plots (Fig. 2), only in the social paradigms were rates reduced in the interval following the first presentation (Supplemental Fig. 5a- c, right graphs).

In a variety of cognitive domains that do not intrinsically involve stimulus repetition (*e.g.*, attention, motion discrimination), better behavioral performance or prediction has been linked with the activity rate of stimulus-tuned cells^32^. Moreover, it is postulated that participants of a neuronal assembly will show increased activity together^33^.

In search of a possible neuronal assembly with increased activity, we created a second threshold of activity. Neuronal activity in trials, when the stimulus was in the cage, was compared to their baseline activity. Neurons that showed over a two standard deviation increase were considered to be “greatly-active” in the trial. From the 744 neurons collected in the SHD test, 197 greatly increased their activity in the first presentation of the conspecific (Fig. 3a, orange). The fraction of greatly-active neurons remained consistent across the following repetitions of the conspecific (T2-T5), at an average of 41%. Similarly, from the 266 active neurons found in the first presentation of the object, 105 were greatly-active (Fig. 3b, orange) with a stable average of 30% in the following habituation trials. Finally, from the 262 active neurons identified in the first presentation of the conspecific in the SRD test, 104 were greatly-active (Fig. 3c, orange) with a stable average of 35% in the following recognition trials.

**Figure 3.**
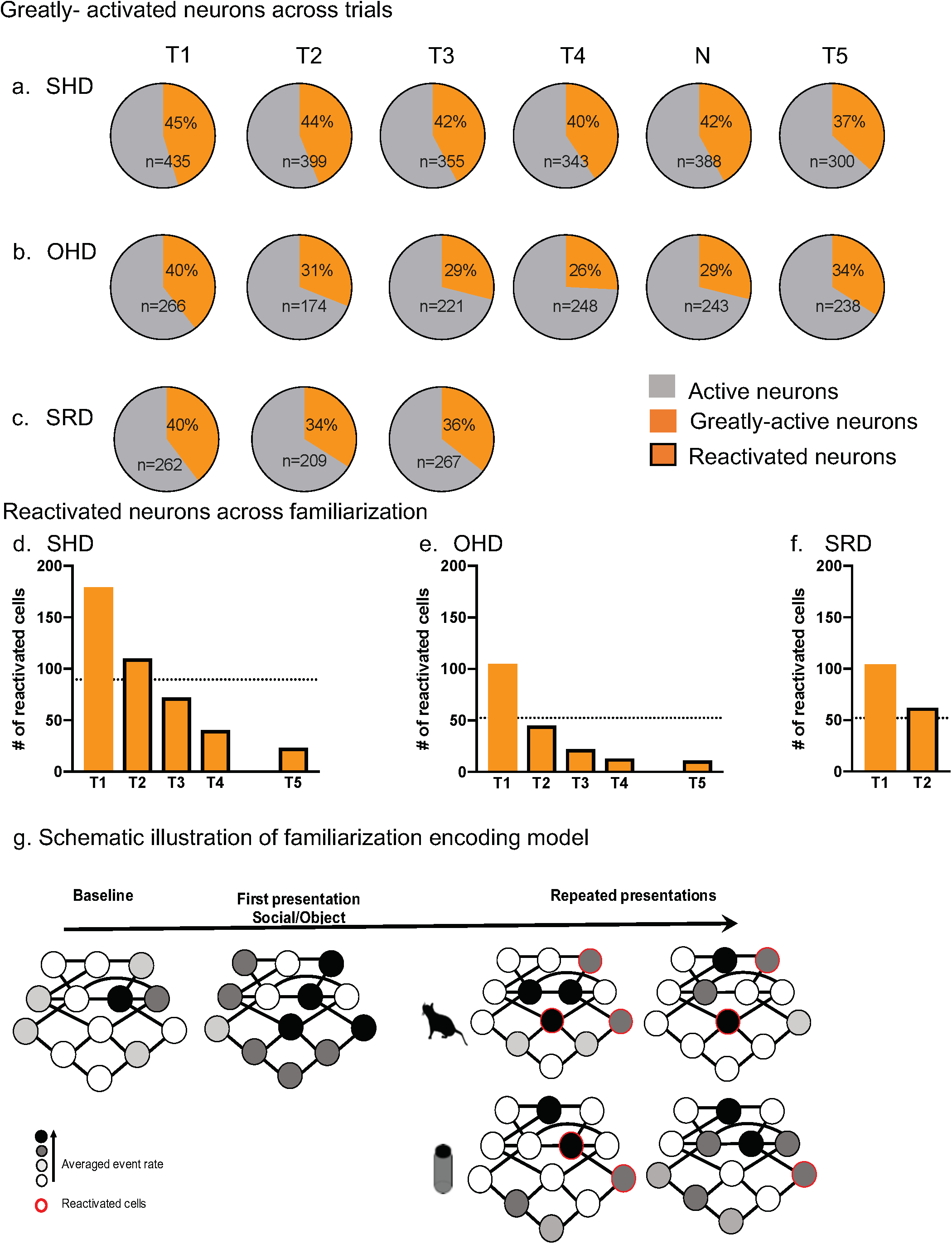
Greatly-active neurons across trials and ensemble reactivation. **a-c.** The fraction of the greatly-active (orange) neurons from the active (gray) neurons across trials was similar in all three paradigms (n = the total number of neurons in trial). **d-f**. Number of reactivated neurons identified as greatly-active in T1 along familiarization (trials T2-T5). **d.** Over 50% of the greatly-active neurons identified in T1 of the SHD paradigm were reactivated in T2 and the number declined throughout familiarization (T3- T5). **e.** Less than 50% of the greatly-active neurons identified in T1 of the OHD paradigm were reactivated in T2 and the number declined throughout familiarization (T3-T5). **f.** Over 50% of the greatly-active neurons in T1 of the SRD paradigm, were reactivated in T2. **g.** Schematic illustration of the possible mechanism underlying encoding of social or inanimate stimuli familiarization.

Taken together, these results demonstrate that while a subset of pyramidal neurons in dCA2 respond to repetitions of both the social and inanimate stimuli with greatly increased activation, stimulus-dependent gradual suppression was only observed in the process of social familiarization.

### Potential reactivated ensembles

To further characterize the neuronal changes possibly underlying familiarization with the conspecific, we aimed to locate a physical substrate of memory. Since real-time memory representation is often achieved by coordinated activity of a neuronal ensemble^34^, we limited the next analysis to potentially selectively responding neurons in the first presentation of the stimulus, the greatly-active neurons (Fig. 3 d-f, T1). We then followed the reactivation of these neurons along the familiarization process. We identified a subset of neurons that were reactivated along all trials of familiarization as well as in the recall trial that followed the interference of a novel conspecific and the 1-hour interval (Fig. 3d, T2-T5). Similarly, a subset of neurons was reactivated in the recall of the inanimate object (Fig. 3e, T2-T5). While in both social and object paradigms (SHD, OHD) we observed a decreased number of reactivated neurons across familiarization, the decline was greater for the object paradigm. Moreover, in both social paradigms (SHD, SRD), over 50% of the greatly-activated neurons identified in the first presentation were reactivated in the second trial although the intervals between the two were very different: 5 min in the SHD paradigm (Fig. 3e; T2), and 30 min in the SRD paradigm (Fig. 3f; T2). Taken together, we suggest the number of reactivated neurons is stimulus dependent. Previously, we showed that CA2 can increase the salience of social signal, through activation of Avpr1b^4^. Our current data indicate that pyramidal neurons of rostral dCA2 may encode different stimuli by utilizing different mechanisms. We have included a possible model to describe the distinct coding scheme for social versus object inputs (Fig. 3g and discussed below).

## Discussion

Familiarity among individuals is viewed as a principal variable in the facilitation of social behavior and its formation thus underlies social cognitive abilities. Current literature supports the essential role of the mouse CA2 in social memory formation^3, 4, 35^, as well as the region’s pyramidal neurons reliance on synaptic potentiation through activation of the Avpr1b^4, 7^. Moreover, the unique functions and characteristics of the CA2 place it in a highly strategic position to be the local integrator of various spatial, temporal, object and social inputs to facilitate or enable memory acquisition^3-5, 36^. The multiple inputs together with its intra- and extra-hippocampal connections, may yield a multi-dimensional “cognitive map” whose creation relies on parallel processing of different cues. Still, studies describing dCA2 neuronal activity in response to novel object or social presentation are inconsistent^5, 29^.

Our experimental approach allowed us to extend previous studies and demonstrate that while dCA2 pyramidal neurons are responsive to both social and inanimate stimuli, distinct neuronal coding schemes are utilized for the short-term processing of the two. A limitation to the study could arise from the possible neuronal responses to other associated features of the task, including spatial movement, immobility and time. First, we did not observe changes in immobility along the trials within each paradigm. Moreover, the unique population of N-units described in a recent study^31^ demonstrated increased activity during immobility whereas we observed a decreased activity along trials of familiarization. Second, CA2 pyramidal neurons show consistent activity patterns along rats’ consecutive visits to the same shape box^36^. Also in that study, changes in CA2 activity were reported to occur over extended periods of time (hours to days). Thus, while many or all inputs may converge at the CA2, given the unchanging environment across our stimuli repetitions, the lack of changes in immobility, and the overall short intervals between trials, we argue that the transient changes we observed were stimulus-dependent and may be viewed as recognition encoding.

Studies have shown that the first presentation of an object or face evokes a distributed pattern of activity in the relevant neural population in which sparsity and selectivity are seen upon familiarization^27, 28^. Sparseness, which has been described in the dorsal hippocampus in general^30^, in addition to having metabolic and efficiency advantages, allows compacted coding of memories^37^, increased input dimensionality, facilitated learning, and reduced noise^38^. Thus, our findings extend the understanding of the neural mechanism of learning to the CA2 with its role in forming social memories performed by its pyramidal neuronal population that also express the CA2 marker, the vasopressin 1b receptor.

When we repeatedly encounter a stimulus in the environment, we become faster and more accurate at identifying it. Improved recognition memory is often attributed to region-specific mechanisms of repetition priming and suppression which reflect improved and efficient encoding that apply over a range of repetition lags^9, 39, 40^. Both priming and suppression occur under the same experimental conditions and often may be associated^41^ and potentially contribute to the formation of long-term representation of stimuli repeatedly presented. Here, we reveal that neuronal activity recapitulates sniffing behavior only toward the conspecific and not the inanimate object.

The suppression may reflect an increased threshold of the pyramidal neurons whose excitability is known to be regulated by the Avpr1b^7^ as well as by the recruitment of interneurons, possibly basket cells, previously shown to have many connections with pyramidal neurons within the local circuitry of CA2^42^ and gate their output^43^. The shifts in excitatory-inhibitory balance could allow CA2 to operate as a comparator when processing multimodal sensory inputs. Nevertheless, invariance may be seen in regard to some stimulus dimensions^39^ which could explain the lack of a significant increase in the calcium event rate in the dishabituation trial when a novel (N) stimulus is presented in the SHD test. The suppression, together with the stable percentage of greatly-active neurons across repetitions, could be viewed as a mechanism in which the greatly-active neurons hold the relevant stimulus-specific information creating a “sharpened” representation of the social cue but also one that facilitates novelty detection.

A major step in understanding how social memories are encoded in CA2 includes locating possible neuronal substrates of these memories. In the hippocampus, coding of an episodic representation may involve both neurons that encode the particular configuration of the stimuli and neurons that encode features that are common among episodes^44^. Our data demonstrate the possible generation of a memory trace with the reactivation of a subset of neurons throughout the recognition of both the social and non-social stimuli. Still, more neurons were recruited during the process of social recognition.

Human studies demonstrate correlation between hippocampal mechanisms and the building of conjunctive representation^45^. Using blood oxygenation level-dependent contrast, familiar items have been shown to elicit smaller hemodynamic responses in human anterior medial temporal lobe^46^. High resolution magnetic resonance imaging studies suggest declarative memory is processed differentially depend on the specific hippocampal subregions, with evidence of CA2/3 contribution to associative encoding^47^. Importantly, cognitive dysfunction is associated with poor psychosocial functions and is a highly relevant dimension of psychiatric disorders^48^. Still, the neuronal mechanisms underlying it remain mostly elusive. Our work uncovers distinct neuronal encoding schemes in pyramidal neurons in dCA2 for social versus inanimate object recognition memory. These coding schemes could allow the CA2 to orchestrate the processing of the multimodal inputs it responds to, including social and inanimate objects, space and time. This would enable the appropriate convergence of inputs as well as the facilitation of their conveyance within the neural circuits. This deeper understanding may also serve as a framework for modeling predictive coding of social memories in CA2 and within the hippocampal microcircuit as well as promote discoveries of the fundamentals of episodic memory architecture and possible interventions. In the future we aim to extend our understanding of the specific role of the Avpr1b in maintaining the neuronal activity scheme during social memory encoding and gain more insights into the neuronal correlates of social memory.

## Acknowledgements

The authors wish to thank Drs. Harold Gainer, Mark Histed and Barry Richmond for reading early drafts of the manuscript and Bruno Averbeck for helpful discussions. We thank Dr. Sun Jung Kang for statistical advice. Emily Shepard provided excellent technical help, including colony maintenance and genotyping. Dr. Kuan Wang provided parts of the MATLAB code used for analysis. We also thank the animal care staff for excellent husbandry. Heon-Jin Lee is currently supported by the National Research Foundation of Korea (MSIT 2017R1A5A2015391)

