## Supplemental information for "Recognition memory via repetition suppression in mouse hippocampal dorsal CA2 pyramidal neurons expressing the vasopressin 1b receptor"

#### Supplementary information

#### Materials and Methods

##### Animals

All housing and procedures were conducted according to US National Institutes of Health guidelines for animal research and were approved by the National Institute of Mental Health Animal Care and Use Committee. Mice were maintained on a 12-h light cycle (lights off at 1500h) with *ad libitum* access to food and water. Adult Avpr1b$^{+/Cre}$ male mice were used for experiments, and singly housed (cage dimension:7.5x7x12.5 inches, WxHxL) following surgery. Ovariectomized (OVX) BalbC female mice (obtained from Jackson Laboratories, Bar Harbor, ME, USA, at 8 weeks of age) were used as social stimuli and singly housed between experiments.

##### *In situ* hybridization histochemistry of neural transcripts

Mice were anesthetized with isoflurane, decapitated and the brains fresh frozen on powdered dry ice. Brains were sliced on a cryostat and 16µm sections were collected onto slides and kept frozen until the time of assay. Sections were used for both methods for *in situ* hybridization histochemistry: Affymetrix ViewRNA [1] or Hairpin Chain Reaction [2]. Details about the probes are presented in Supplementary Table 1.

##### *In situ* hybridization using the ViewRNA method

Tissues were prepared as described above. Next, using ViewRNA [1] reagents (Duplex kit; Catalog number: QVT0013; Santa Clara, CA), we prepared working probe set solutions (types 1 and 6; Fast Red and Fast Blue, respectively) at a 1:40 dilution in prewarmed probe set diluent. The probe solutions were applied to the sections and hybridized for 2 hours at 40°C. The sections were next washed three times for 2 min in wash buffer with constant and vigorous agitation and then placed in storage buffer overnight. Next, signal amplification and detection were performed with kit reagents: The Preamplifier Mix QT (25 min, 40°C), Amplifier Mix QT (15 min, 40°C) with wash steps in between. Label Probe 1-AP was then added at a dilution of 1:1000 in prewarmed label probe diluent QF for 15 min at

40°C. For development, the AP Enhancer Solution was added for 5 min at RT, and then immediately the Fast Red substrate was applied for 30 min at 40°C. After the development of the first probe, the reaction was stopped by first washing the sections in wash buffer, and then adding AP stop QT buffer for 45 min at RT in the dark. Label probe 6-AP was then added at a 1:1000 dilution in prewarmed label probe diluent QF, for 15 min at 40°C. A Fast Blue substrate was applied and sections were incubated for 30 min in the dark at RT. Next, sections were rinsed briefly in PBS and then in the wash buffer. Finally, the tissues were briefly washed in PBS and counterstained with DAPI in 0.1M Tris-HCl, pH 8, for 1 min.

***In situ* hybridization using the Hairpin Chain Reaction method**

We used the Hairpin Chain Reaction (HCR) [2], with some modifications, in our mapping study to locate transcripts for Avpr1b and the CA3 marker, Bcl-2-related ovarian killer protein (Bok). Slide-mounted, fresh-frozen sections were fixed in 4% formaldehyde/PBS at room temperature (RT) for 5 min. Following fixation, sections were briefly washed in PBS at RT twice for 1 min each. Then the sections were incubated in a solution of acetic anhydride in triethanolamine, pH 8, for 10 min. The sections were first processed through a series of ethanol steps (70%, 1 min; 80%, 1 min; 95%, 2 min; 100%, 1 min) followed by $CHCl_3$ for 5 min and then back through ethanol (100%, 1 min; 95%, 1 min). Then the sections were air dried. Next, probes (at a working concentration of 4.0nM) were added to a nucleic acid mix (100 µg/ml salmon sperm DNA, 250 µg/ml yeast total RNA, 250 µg/ml yeast tRNA; all Sigma-Aldrich), heated to 65°C for 5 min, and then cooled on ice for 5 min. This mixture was added to the hybridization buffer [50% formamide/600 mM NaCl/80 mM Tris-HCl, pH 7.5/4 mM EDTA/0.1% sodium pyrophosphate/0.2% SDS/0.2 mg/ml sodium heparin/2% sodium polyacrylate]. The probe cocktail was added to the sections and then incubated in a humidified chamber for 24 hours at 37°C. The next day, sections were washed in 1xSSPE (150 mM NaCl, 10 mM $NaH_2PO_4$, and 1 mM EDTA, pH 7.4) four times for 30 min each at 37°C with gentle rotation. The sections were then washed in 1x SSPE for 5 min at RT and then in 5xSSPE for five min. The hairpin amplification took place with minimal light exposure, including dimming lights when possible. At a working concentration of 60 nM, each hairpin was heated separately at 95°C for 1.5 min and then cooled to RT for 30 min. The sections were then hybridized at RT for 24 hours. The next day, sections were washed at RT in 5xSSPE with 0.1% Tween-20 four times for 30 min each with gentle rotation. The sections were briefly rinsed in 5xSSPE and then counterstained with DAPI in 5xSSPE for 1 min. In order to minimize any autofluorescence, sections were then incubated in 1x True Black in 70% ethanol for 2 min. Finally, the sections were washed at RT in PBS 3 times for 5 min each, followed by a quick dip in 70% EtOH before air-drying.

**Imaging and analysis**

All initial images were taken on a Nikon Eclipse 50i binocular microscope with a CoolLED fluorescence source (Andover, UK). The Fast Red (type 1) and HCR Alexa fluor 594 developed probes were visualized using a TRITC filter set (EX: 510-550, EM: 590-610). The Fast Blue (type 6) and Alexa fluor 647 developed probes were visualized using a far red filter set (EX: 610-640, EM: 750-810). Alexa fluors 488 and 546 developed probes were visualized using green (EX: 450-490, EM: 500-550) and orange (EX: 538-562, EM: 570-640) filter sets, respectively. DAPI was visualized with a blue filter set (EX: 325-375, EM: 435-485). Images were captured using a QImaging Retiga 4000R digital camera and imaging software from BioVision Technologies (iVision, Exton, PA, USA). Lower resolution scans were also obtained using a Zeiss Axio ScanZ1 (20x objective) and ZENlite software (Thornwood, NY, USA).

**Behavioral paradigms**

Tests were performed in the light cycle (lights off at 1500h) in a dim testing room adjacent to the mouse housing room. All interactions were videotaped and coded offline by a trained observer blind to the genotype of the test mice using Noldus Observer XT 12 (Noldus Information Technology, Leesburg, VA). Prior to testing, mice were allowed to habituate to a neutral cage for 30 min in the testing room. For social habituation-dishabituation (SHD, Supplemental Fig. 3a), an unfamiliar OVX female was introduced repeatedly for 2 min with 5-min inter-trial intervals (ITI; T1-T4-stim 1), then a novel OVX female was introduced for 2 min (N-stim 2). After a 45-60-min ITI, the original female was reintroduced once again for 2 min (T5-stim 1). During the inter-trial intervals, the female was returned to her home cage while the experimental animal remained in the testing cage. Mice were allowed to fully interact and anogenital and all body sniffing were measured. Object habituation-dishabituation (OHD, Supplemental Fig. 3c) was run similarly with an unfamiliar object (Lego piece, Billund, Denmark, as stim 1) then a novel one (N, 20ml scintillation vial with purple water as stim 2). Sniffing of the object, indicated by head movement and snout within 1 cm of object, was measured. The social recognition-discrimination (SRD, Supplemental Fig. 3e) test was modified from Engelmann et al.[3]. An unfamiliar OVX female was introduced for 5 min (T1-stim 1), then reintroduced after 30-min (T2-recognition; stim 1), and then again after another 30-min together with a novel female (T3/N- discrimination; stim1 and stim 2). Mice were allowed to fully interact and anogenital and all body sniffing were measured. The discrimination score was calculated as (sniffing novel - sniffing familiar)/(sniffing novel + sniffing familiar). Mice were video-tracked in Ethovision (Noldus Information Technology) for immobility periods.

**Adeno-associated viruses (AAV)**

AAV1-Syn-Flex-GCamp6s.WPRE.SV40 (University of Pennsylvania Vector Core, Philadelphia, PA, USA) was stored at -80ºC until used. On day of the experiment, the virus was kept in dry ice until diluted with normal 0.9% saline to 1.8 $x10^{12}$ genome copies/ml.

**Stereotaxic surgery and calcium imaging**

Stereotaxic procedures were performed in accordance with Animal Care and Use Committee of the National Institute of Mental Health. Male mice (10-16 weeks old) were anaesthetized with intraperitoneal 0.1 ml/per 10g body weight ketamine/xylazine (100 mg/16 mg, respectively; per 10 ml saline) and topical lidocaine ointment 5% (Fougera, Melville, NY, USA). After placement on a stereotaxic frame (David Kopf Instruments, Tujunga, CA, USA), the skull was exposed and perforated with a drill to allow virus delivery via a 5 µl microliter syringe (26 g, Hamilton, Reno, NV, USA) and Micro4 micro-syringe pump (World Precision Instruments, Sarasota, FL, USA) (300 nl at 40 nl/min at bregma:

anterior-posterior (AP): -1.06 mm, medial-lateral (ML): 0.6 mm, dorsal-ventral (DV): -1.75 mm). Following >2 weeks of recovery, mice were prepared for calcium imaging (n=5) as described[4]. Briefly, a gradient-refractive lens (outer diameter, 0.5 mm; length ~6.1 mm; numerical aperture=0.5) was implanted directly over the dCA2 pyramidal neurons (AP: -0.94 mm; ML:0.4 mm; DV:-1.78 mm). Following >2 weeks, a baseplate was attached to the skull with dental cement under isoflurane anesthesia (1.5-2.0%). Mice were habituated to the microscope mounting over the following 3 days. All stereotaxic injections and implants were histologically verified (see Supplemental Information). Calcium imaging was acquired with Vista HD 2.0 system (Inscopix, Inc., Palo Alto, CA, USA) at a rate of 20 frames/s, while synchronized with the behavioral recording software (Ethovision). LED power was kept under 15% with increased gain (up to 4). Mice who showed no change in sniffing duration along the repetitions of the paradigm or novel presentation were excluded from further analysis.

Five mice were included in neuronal data analysis: social habituation-dishabituation (mouse 1, mouse 2, mouse 3, mouse 4), object habituation-dishabituation (mouse 1, mouse 4, mouse 5), and social recognition-discrimination (mouse 1, mouse 3, mouse 4). A total of 744 cells were analyzed in the social habituation-dishabituation paradigm (with distribution in each mouse being: 160, 290, 147, 147), 433 cells in the object habituation-dishabituation paradigm (with distribution in each mouse being: 96, 149, 188) and 451 cells in the social recognition discrimination paradigm (with distribution in each being: 154, 150, 147).

Calcium recordings were spatially down sampled (factor of 4) and motion corrected (Image Stabilizer, ImageJ, US National Institutes of Health). The fluorescent traces from individual neurons were extracted by principal component and independent component analysis (PCA-ICA) (Mosaic, Inscopix). This multi-steps analysis, inspired by Mukamel *et al.* [5], allows the identification of signals from independent components (neural soma) in acquired movie data. After visually inspecting the spatial filter together with the temporal signal, data was further analyzed with a custom-written MATLAB script (available upon request). Based on the sensitivity and kinetics of GCaMP6s [6], we detected $Ca^{2+}$

events in the identified neurons with a threshold of peak dF/F0>5 standard deviations above the median absolute deviation. As described previously [7], we identified the rising phase of each calcium event and calculated the first derivative with a 200ms moving window. Calcium events were binned over the 2 min (either an exposure trial, end of baseline, or inter trial interval) for every identified neuron (event rate). Neurons were considered active if they responded with at least one $Ca^{2+}$ event in any designated bin of 2 min (either an exposure trial, end of baseline, or inter trial interval). Analysis for comparison of calcium events across trials included all identified neurons, including silent ones in the trial. Percent of active neurons in trial was calculated as $\frac{number\ of\ active\ neurons}{total\ number\ of\ neurons} \times 100$. While the data is non-normally distributed and analyzed with non-parametric statistical tests, for simplicity, bar graphs present the mean with SEM.

Neurons were considered greatly-active if their calcium event rate during any 2 min trial/inter-trial interval was at least 2 standard deviations greater than their mean calcium event rate during the 8-10 min of baseline. All identified greatly-active neurons were then pooled for further analysis including comparison of percentage of greatly-active neurons and reactivation. Percentage of greatly-active was calculated by $\frac{number\ of\ greatly-active\ neurons}{number\ of\ active\ neurons} \times 100$. In the first trial of any paradigm (T1), neurons meeting either active or greatly-active criteria were considered new if they were silent at baseline. For reactivation analysis, the greatly-active neurons identified at the first presentation (T1) were followed along the repetitions.

In order to determine whether the pattern of neuronal activity was reliably different between the trials with the stimulus and the inter-trial intervals (ITI, stimulus is absent), we deconvolved neuronal events and aggregated them into counts over 500 msec bins (*i.e.,* each 2 min trial was of comprised 240 bins). For a given animal, we identified all bins in any trial or ITI where at least one neuron fired and aggregated them into a single dataset. Each of these bins was labelled into two classes, reflecting whether it came from a trial or an ITI. The exact number of bins varied across experiments (SHD, OHD, SRD) and animals, but there were more than 500 of each in almost all instances. The number was not consistently larger for trial or ITI.  Given the data, we used a linear support vector machine classifier (with default parameter setting 1) to distinguish whether a bin came from a trial or an ITI, given a vector with the aggregated counts of activity during that bin for all neurons. We carried out a balanced cross-validation procedure over all time bins. In this procedure, a subset of bins was left out in turn, the classifier was trained on the other bins, and applied to the test bins to generate predictions. Classifier performance was measured as the fraction of bins where the classifier predicted correctly whether the bin came from a trial or ITI. For different animals and experiments there was generally more of one type of bin than the other, therefore we balanced each dataset by sampling the most frequent type to be as numerous as the least frequent one. The cross-validation procedure was then run on a balanced dataset. Because this introduces stochasticity in the results, we repeated the entire analysis 100 times and report the average accuracy result across repeats. Furthermore, we took into account the possibility of temporal dependence between consecutive bins and left out groups of adjacent bins together to make each test set. In this manner, no training set bins were directly adjacent to test set bins.  Accuracy results were tested using a binomial test with a null hypothesis of classifier chance performance (accuracy=0.5) [8]. We corrected for multiple comparisons (using Bonferroni correction) to take into account the number of animals across the three experiments. We also compensated for the possibility that prediction in adjacent time bins are correlated, by using half of the effective number of bins in testing. Null hypothesis that trial and ITI neuronal activity is not distinguishable could be rejected with significance at the corrected p=0.01 level.

In order to determine whether the pattern of neuronal activity contained information about sniffing, we deconvolved neuronal events and aggregated them into counts over 500 msec bins. Bins were labelled into two classes, reflecting the presence or absence of sniffing at that point. In general, there were more bins without sniffing than bins where it took place, but the exact ratio varied across experiment (SHD, OHD, SRD), animal, and trial. Given this data, we used a linear support vector machine classifier (with default parameter setting 1) to predict whether sniffing was taking place in a time bin, given a vector with the aggregated counts of activity during that bin for all the neurons. We carried out a leave-one-out cross-validation procedure over all the time bins within a trial. In this procedure, each bin was left out in turn, the classifier was trained on the other bins, and applied to the test bin to generate a prediction. Classifier performance was measured as the fraction of test bins where the classifier predicted correctly whether sniffing was taking place. There are, in general, fewer bins where sniffing occurs than ones where it does not. To prevent the classifier from doing well by predicting no sniffing by default, we balanced each dataset set by sampling no-sniffing bins, so that they were as numerous as sniffing bins. The cross-validation procedure was then run on a balanced dataset. Because this introduces stochasticity in the results, we repeated the entire analysis 100 times and report the average accuracy result across repeats. Accuracy results were tested using a binomial test with a null hypothesis of classifier chance performance (accuracy=0.5) [8]. We corrected for multiple comparisons to take into account the number of animals across three experiments, and the number of trials per animal, using Bonferroni correction.

**Statistics**

Statistical analyses were performed using Prism (version 7, GraphPad Software, San Diego, CA, USA) and expressed as mean ± sem. For behavioral data either repeated measures ANOVA or unpaired t-tests were used with Tukey's HSD post hoc comparisons. For imaged mice sniffing, repeated measures one-way ANOVA with test for linear trend across trials or Mann-Whitney test were used. For analysis of immobility, 2-way ANOVA was used for comparing social versus object habituation trials. A Kruskal-Wallis test or Friedman test was used for comparisons across paradigms and within the social recognition one. Neural data was tested with D'Aostino-Pearson omnibus for normality, followed by a non-parametric Friedman test and Dunn's multiple comparison tests. Comparisons between trials with and without stimuli were performed with Wilcoxon pairs signed rank test with the Pratt method. Comparisons between social and object trials were performed in SAS (SAS Institute, Cary, NC, USA) with generalized linear mixed model for non-normal data. The number of active or greatly-active neurons was tested with Fisher's exact test. #, $P$ <0.0001; ***, $P$ <0.001; **, $P$ < 0.01; *, $P$ <0.05.

**Code availability.**

The MATLAB code for analysis is available from the corresponding author upon request.

#### Results

##### Avpr1b is a CA2 marker for pyramidal cells

Supplemental Fig. 1 shows that transcripts for Avpr1b and the CA3 marker Bok are not colocalized in the same neurons. Furthermore, those neurons expressing Avpr1b co-express the dCA2 markers PCP4, Map3k15 and Amigo2. Fig. 2a shows that heterozygous Cre recombinase knock-in mice co-express Avpr1b and cre recombinase in the same neurons, as previously described [9]. Finally, 87% of non-glial cells in the CA2 express Avpr1b (Fig. 2b-e). The remaining non-Avpr1b neurons express the interneuron marker glutamic acid decarboxylase [10].

##### Behavioral validation of transgenic mouse model

In our KI line, the Avpr1b gene is replaced with the Cre gene, so homozygous KI mice lack Avpr1b gene expression entirely (Avpr1b$^{Cre/Cre}$). In agreement with our earlier knock-out mice [11], Avpr1b$^{Cre/Cre}$ mice performed well in a social habituation-dishabituation (SHD) paradigm. In a series of 2-min exposure trials (T1-T4 with stimulus mouse 1; N with stimulus mouse 2) separated by 5-min inter-trial intervals (ITI) (Supplementary Fig. 3a), mice demonstrated decreased sniffing duration upon familiarization to the stimulus and a significant increased investigation of the novel stimulus. Moreover, Avpr1b$^{Cre/Cre}$ mice showed decreased sniffing of the already familiar stimulus (T5 with stimulus mouse 1) when presented again following a longer (~1 hour) interval (Supplementary Fig. 3b). Some aspects ("what" and "when" but not "where") of object recognition are weaker in the Avpr1b knock-outs [12]. In our current paradigm no differences arose in the Avpr1b$^{Cre/Cre}$ mice in the object habituation-dishabituation (OHD) paradigm (T1-T4 with stimulus object 1; N with stimulus object 2; Supplementary Fig. 3c). Mice decreased sniffing duration upon familiarization, followed by increased investigation of the novel object

(Supplementary Fig. 3d). In a more challenging social recognition-discrimination (SRD) test (Supplementary Fig. 3e), mice were given two 5-min exposure trials to the same stimulus female with an ITI of 30 min to test for recognition. An additional exposure, following a second 30-min interval, included both the familiar and a novel stimulus female, requiring the experimental mouse to actively discriminate between the two. When comparing the recognition trials (Supplementary Fig. 3f), the WT spent significantly less time sniffing the familiar female in the second trial whereas, as expected, the Avpr1b$^{Cre/Cre}$ mice did not show this decrease in sniffing, demonstrating that inactivating the Avpr1b gene by knocking in the Cre gene results in social memory deficits as previously described when knocking out the Avpr1b [11]. We previously reported deficits in sociability and preference for social novelty in Avpr1b knock-outs [12]. We did not observe any indication of difference between mice in the initial motivation to investigate the social stimulus in both paradigms. Sociablity is not affected by silencing of dCA2 pyramidal neurons [13]. We did not observe a significant difference in the discrimination performance of the two genotypes (Supplementary Fig. 3g), suggesting that by this point, the Avpr1b$^{Cre/Cre}$ mice are able to discriminate between conspecifics. Possible explanations for the discrepancy with the reduced social novelty preference described by DeVito et al.[12], may arise from critical differences in the paradigm design. In the prior study, mice were tested in a 3-compartment apparatus divided by thin plastic mesh with a male stimulus, whereas in the current study mice were tested in an apparatus identical to their home cage that allowed full interaction with an ovariectomized female. Restricted social interactions have been suggested to alter mouse behaviors [14], thus possibly with the design of the current study, Avpr1b$^{Cre/Cre}$ mice are provided with conditions in which discrimination is possible.

Supplementary Table 1

| Probe Name | Symbol | Process | Transcript Target | Probe Pairs | Catalog # |
|---|---|---|---|---|---|
| Vasopressin 1b receptor | Avpr1b | HCR | NM_011924.2 | 30 | not used |
| Vasopressin 1b receptor | Avpr1b | ViewRNA | NM_011924.2 | 20 | VB1-16301, VB6-16894 |
| Adhesion molecule with Ig like domain 2 | Amigo2 | ViewRNA | NM_178114 | 20 | VB6-19241 |
| Bcl-2-related ovarian killer protein | Bok | HCR | NM_016778 | 20 | not used |
| Bcl-2-related ovarian killer protein | Bok | ViewRNA | NM_016778 | 20 | VB1-3033354 |
| Cre recombinase | Cre | ViewRNA | HQ335171 | 17 | VF1-13057, VF6-16306 |
| Mitogen-Activated Protein Kinase Kinase Kinase 15 | Map3k15 | ViewRNA | NM_001163085 | 20 | VB1-20765 |
| Purkinje Cell Protein 4 | Pcp4 | ViewRNA | NM_008791 | 14 | VB1-19888, VB6-19889 |

### Supplementary Figures

#### Supplementary Figure 1

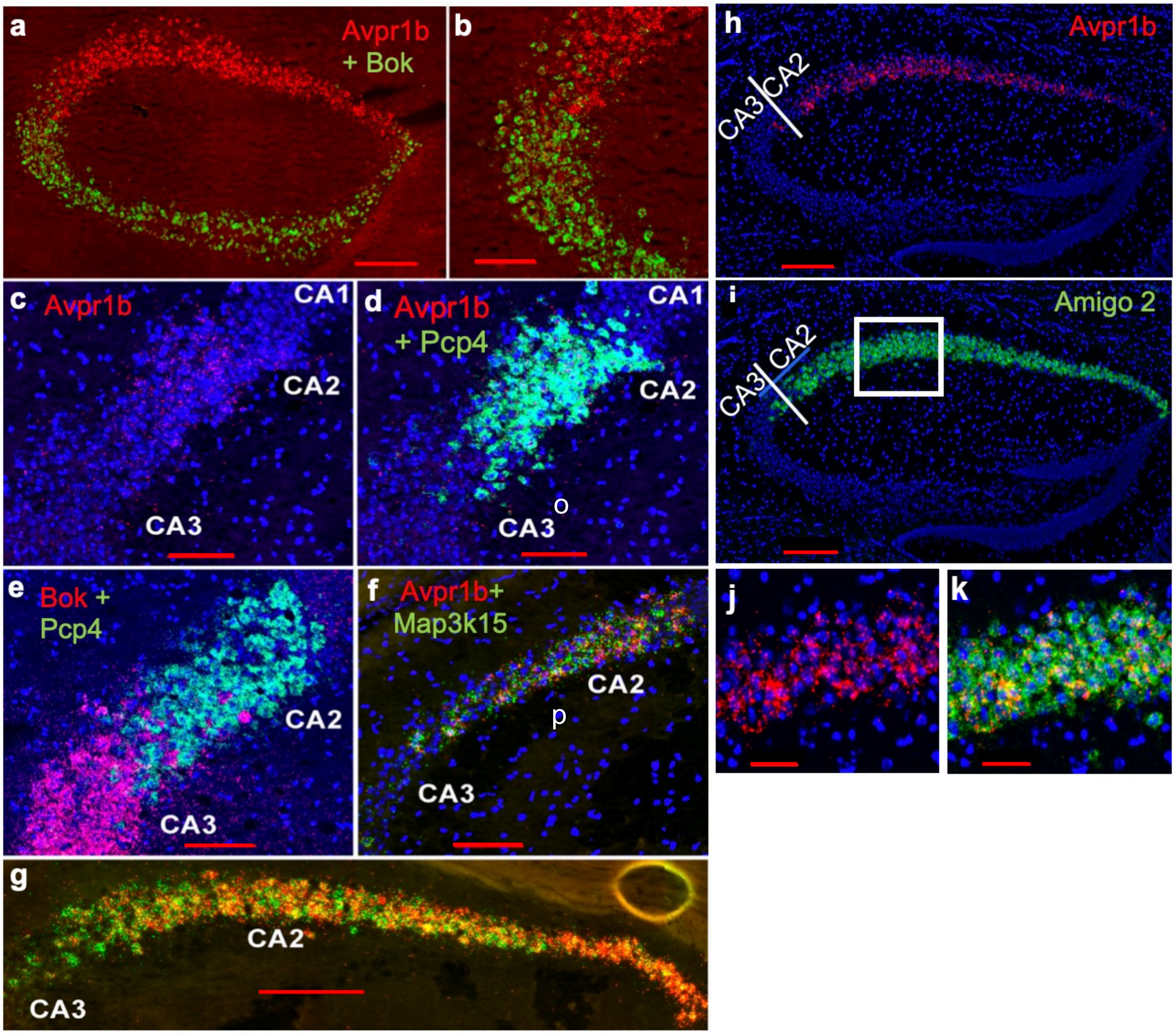

**Supplementary Figure 1**

**Avpr1b as a marker for CA2 hippocampal field.** Panels **a** and **b** show that expression of Avpr1b (red) and the CA3 marker Bok (green) have different distributions whose neurons interdigitate at the inexact CA2/CA3 boundary. Panels **c** through **f** show CA2/CA3 boundaries at a more lateral and caudal level. Panel **c** shows expression of Avpr1b (red) in the CA2 and CA2/CA3 boundary that is colocalized with expression of the CA2 marker Pcp4 in panel **d**. Panel **e** shows the same interdigitation of neurons at the CA2/CA3 border expressing Bok (red) and Pcp4 (green). Panels **f** and **g** show the colocalization transcripts for Avpr1b (red) and the CA2 marker Map3k15 (green) in the dCA2. Panel **f** shows again the imprecise border between CA2 and CA3. Panel **g** is at the level of the lens implant. Panels **h** and **j** show the expression of Avpr1b in dorsal CA2. Panels **i** and **k** show the expression of CA2 marker Amigo2 in the dorsal CA2. Panel **k** shows the colocalization of Avpr1b and Amigo2 in the dorsal CA2. Panels **a** and **b** were processed for *in situ* hybridization histochemistry by Hairpin Chain Reaction and the rest by ViewRNA. Bars are 50μm for panels j and k, 100μm for panels b-f; and 200μm for panels a, g, h, and i.

#### Supplementary Figure 2

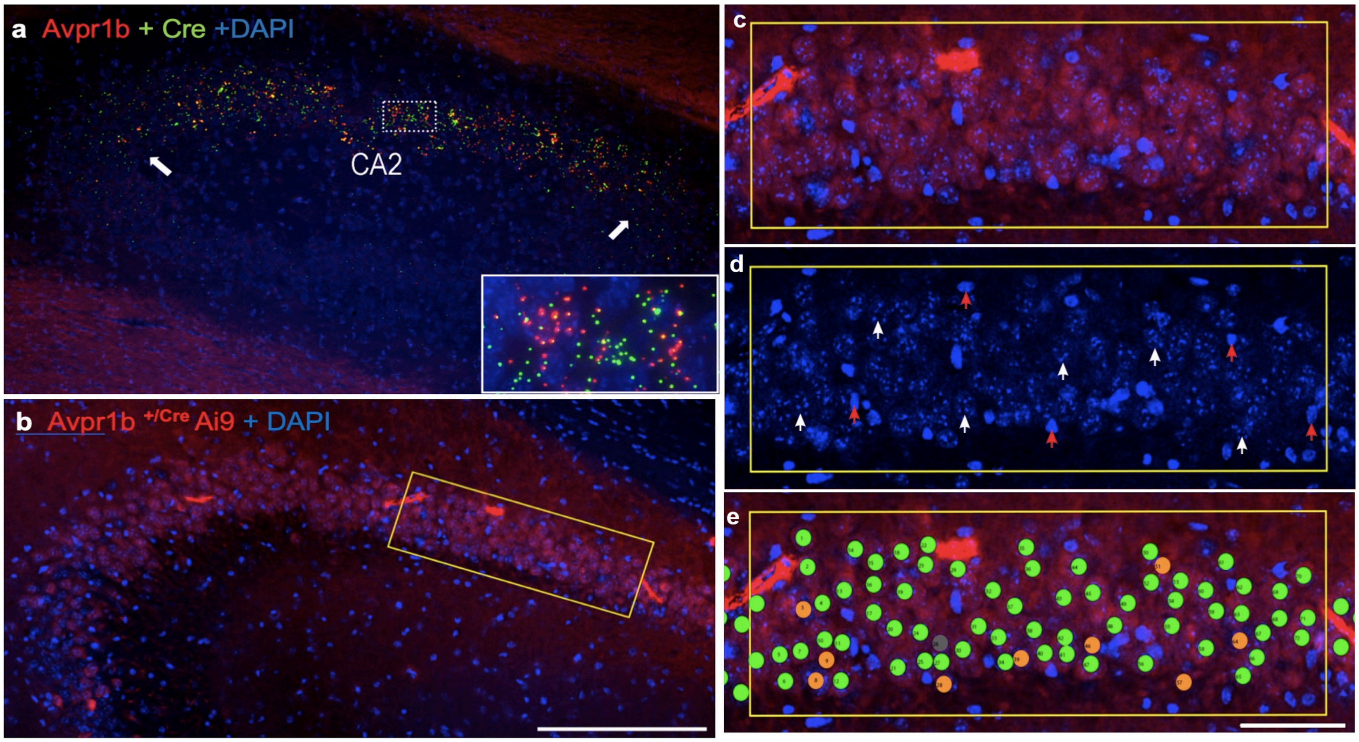

**Supplementary Figure 2**

**Avpr1b is expressed in the vast majority, if not all, of CA2 pyramidal neurons.** Panel **a** is a representative image of double *in situ* hybridization histochemistry on a section (16µm) from a heterozygous mouse (Avpr1b$^{+/Cre}$) demonstrating Cre recombinase (green) mRNA is selectively expressed in neurons that co-express Avpr1b mRNA (red) in dorsal CA2. Arrows indicate the borders of CA2. Panels **b-e** are sections from Avpr1b-Cre Ai9 reporter line at the dorsal CA2 showing expression of tdTomato in Cre-expressing neurons and stained for DAPI. Panel **c** is and area indicated by the box in panel b at higher magnification at which cell counting was performed. The DAPI channel (panel **d**) was used for marker placement and quantification over nuclei. White arrows indicate representative neuronal nuclei on which markers were placed and red arrows indicate compact nuclei, likely glial nuclei, not included in analysis. Panel **e** shows overlaid markers on the DAPI channel in the composite image. Green markers indicate DAPI colocalization with tdTomato (*i.e.*, Avpr1b expression); orange markers indicate no colocalization; a single grey marker indicates a discarded marker because only one tdTomato-positive neuron was seen for two marked nuclei. 87.3% of selected DAPI nuclei were tdTomato-positive. Bars are 200µm in panels a and b and 50µm in the other panels.

**Supplementary Figure 3**

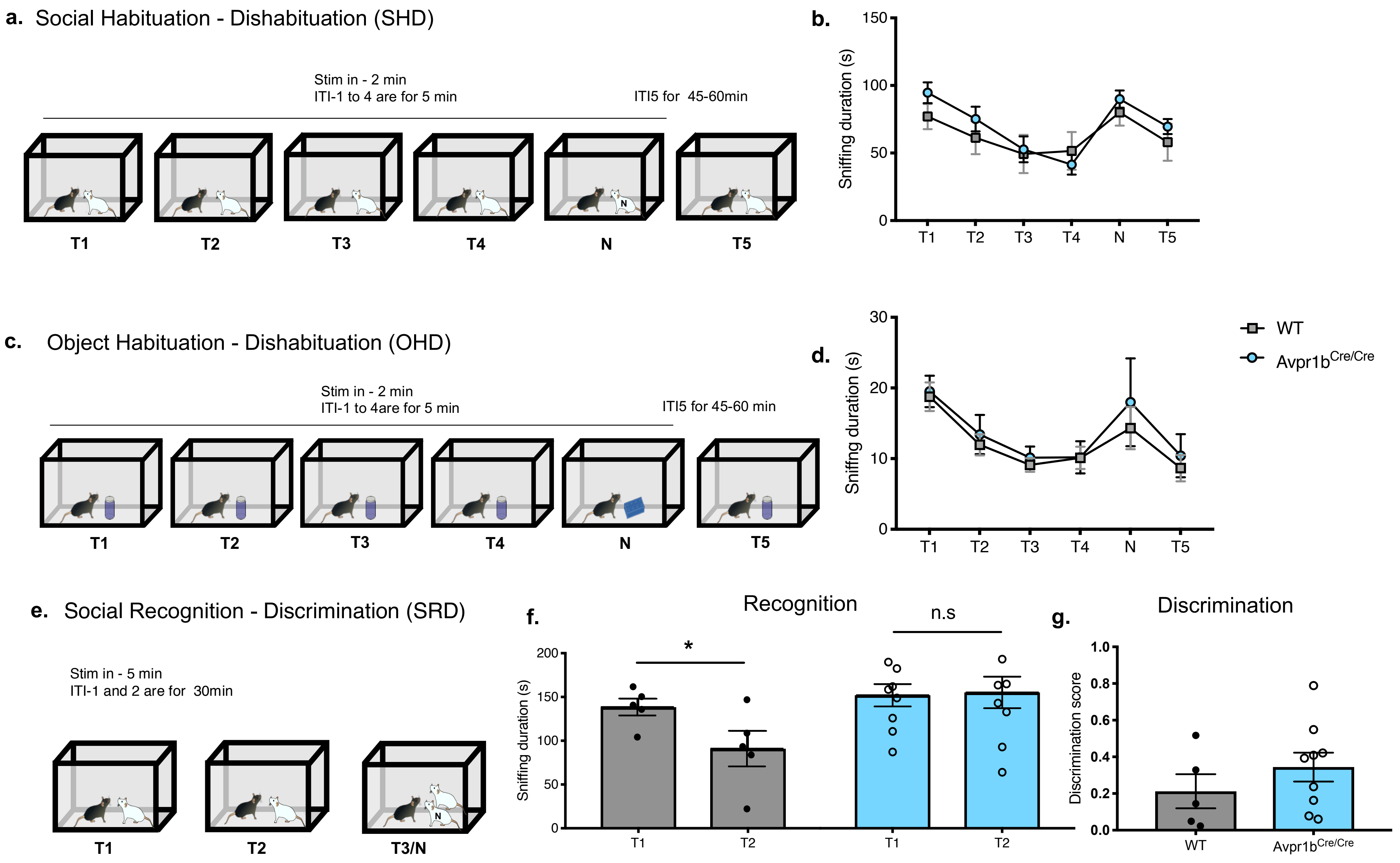

**Supplementary Figure 3**

**Removing the Avpr1b gene results in social memory deficit. a.** Schematic illustration of SHD paradigm (T1-T4: stimulus mse 1; N: stimulus 2 with 'N' annotation; T5: stimulus mouse 1) **b**. Both groups show decreased sniffing duration during habituation (WT=5 mice; Avpr1b$^{Cre/Cre}$=11mice; repeated measures ANOVA genotype X trials: significant effect of trials $F(5,65)=17.82$, $P<0.0001$, with no interaction) followed by increased sniffing of the novel stimulus (within genotype Sidak's multiple comparison $P<0.05$ for both groups). Within genotype comparison show significant decrease in T5 in Avpr1b$^{Cre/Cre}$ mice (Sidak's multiple comparison $P<0.05$). **c**. Schematic illustration of OHD test (T1-T4: lego cube; habituation; N: jar filled with purple water; dishabituation; T5-lego cube again). **d**. Both groups show decreased sniffing duration during habituation (WT=8 mice, Avpr1b$^{Cre/Cre}$=10 mice; repeated measures ANOVA genotype X trials: significant effect of time $F(5,115)=13.65$, $P<0.0001$, with no interaction). Within genotype Sidak's multiple comparisons show for Avpr1b$^{Cre/Cre}$ mice a significant increase in sniffing of the novel stimulus and decreased sniffing of the familiar presented at T5. **e**. Schematic illustration of SRD paradigm (T1-T2: stimulus mouse 1; T3/N: stimulus mouse 1 and 2 with 'N' annotation) **f**. WT group shows decreased sniffing while the Avpr1b-/- group does not (WT= 5 mice, Avpr1b$^{Cre/Cre}$=11 mice; repeated measures ANOVA: genotype X trial: interaction: $F(1,12)=5.179$, $P<0.043$; multiple comparison for trial within genotype: for WT: $P<0.05$ for Avpr1b$^{Cre/Cre}$, $P>0.05$). **g**. Both groups demonstrate similar interest in the novel stimulus presented (unpaired t-test P=0.2902). Data shown as mean±sem.

**Supplementary Figure 4**

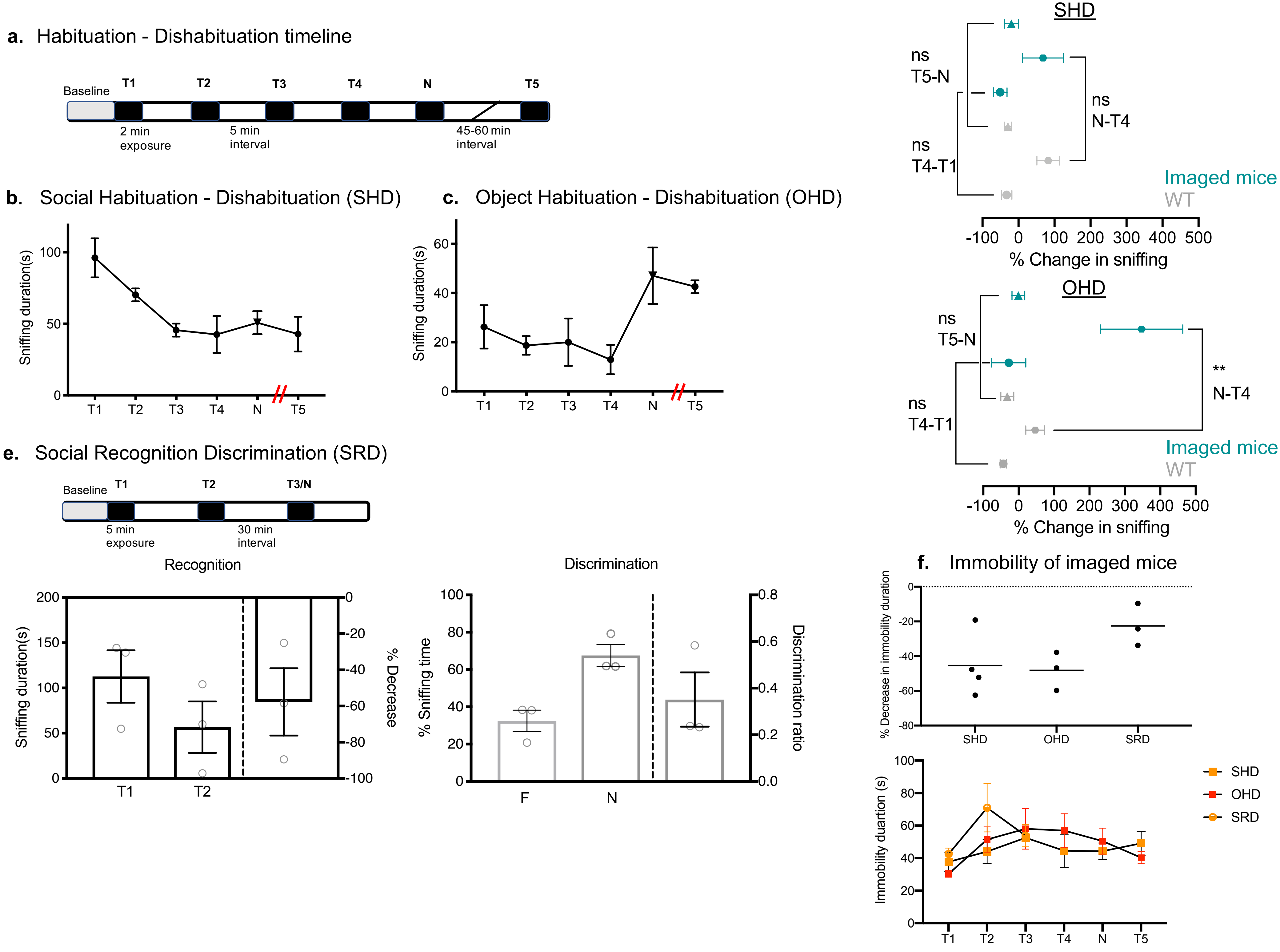

**Supplemental Figure 4**

**Behavioral performance of imaged mice. a.** Timeline illustration for the habituation-dishabituation paradigms. **b.** Sniffing quantification of imaged mice performing social habituation-dishabituation (n=4; line graph) (linear trend slope of -18.54, $F(1,9)=15.83$, $P=0.0032$) **c.** Sniffing quantification of imaged mice performing object habituation-dishabituation (n=3; line graph). **d.** No difference in the percent change in sniffing duration (top graph) between imaged and wildtype mice performing the SHD (Mann-Whitney test: $P>0.1$). Significant difference in the percent change in sniffing for the novel object (lower graph) (Mann-Whitney test: $P>0.03$). e. Timeline illustration and sniffing quantification of social recognition discrimination (n=3). In the recognition trial (T2; left graph), mice showed over a 50% reduction in sniffing time. In the subsequent discrimination trial (right graph), mice showed a preference toward the novel animal presented (N) (Mann-Whitney test $P>0.05$). **f**. All imaged mice decreased their immobility duration in the first trial. No difference was found for immobility duration of imaged mice across trials (SHD vs. OHD Paradigms [2-way ANOVA Trial: $F(2.450,12.25)=2.759$, $P=0.095$; $F(1,5)=0.1325$, $P=0.7307$. There was no interaction]. Comparison of T2 trials or the Novel trials across the three paradigms with Kruskal-Wallis test, $P=0.6$. (data are shown as mean±s.e.m.)

#### Supplementary Figure 5

##### **a**. SHD- Neuronal response in intervals

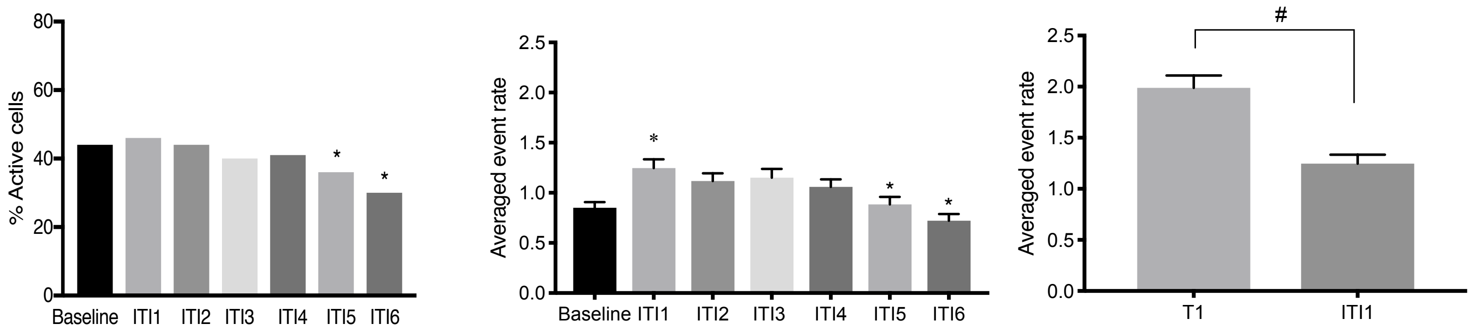


##### **b**. OHD – Neuronal response in intervals

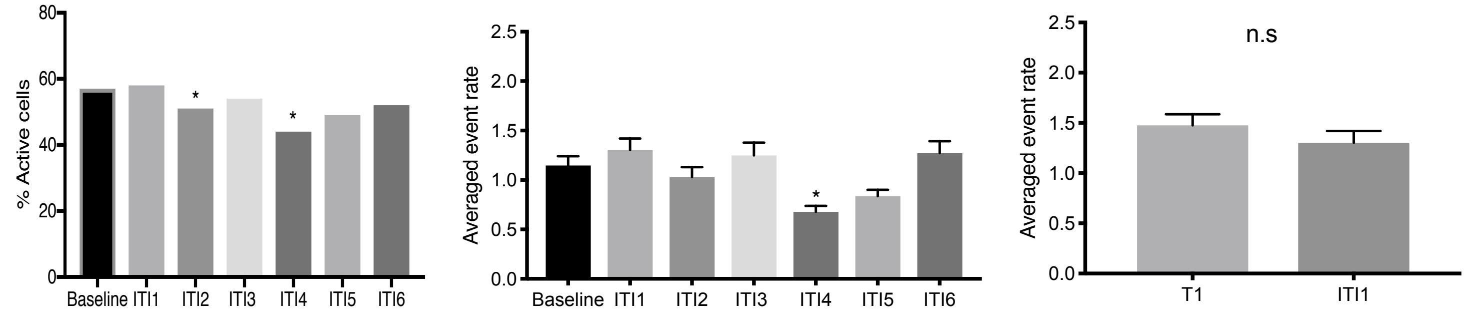


##### **c**. SRD – Neuronal response in intervals

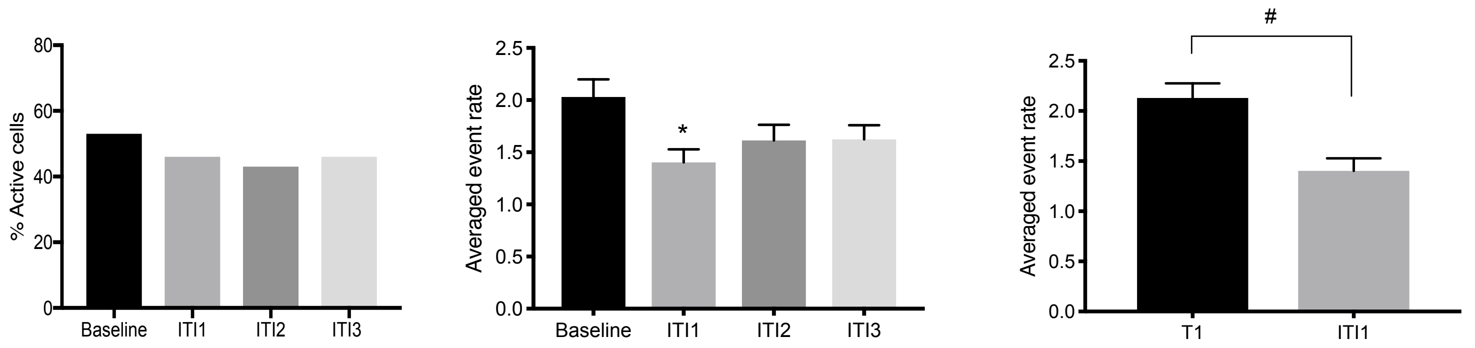

**Supplementary Figure 5**

**Neuronal responses in repeated intervals. a**. (Left graph) SHD: percentage of active neurons across intervals (Fisher's exact test: ITI4-ITI5 *P*<0.05; ITI5-ITI6 *P*<0.03). (Middle graph) Calcium events rate across intervals (n=744 neurons from 4 mice: Wilcoxon signed rank test Baseline-ITI1 *P*<0.003; repeated measures Friedman test: trial X events *P*<0.0001 with Dunn's multiple comparisons test ITI5-ITI1 *P*<0.02; ITI6:ITI1, ITI2, ITI3, ITI4 (Wilcoxon signed test T1-ITI1 *P*<0.0001). **b.** (Left graph) OHD: percentage of active neurons across intervals (Fisher's exact test: ITI1-ITI2 *P*<0.05; ITI3-ITI4 P<0.07). (Middle graph) Calcium events rate across intervals (n=433 neurons from 3 mice; repeated measures Friedman test: trial X events, P< 0.0001; Dunn's multiple comparisons test: ITI4:ITI1, ITI6 *P*<0.03). (Right graph) No change in rate upon removal of the object. **c**. (Left graph) SRD: percentage of active neurons across intervals. (Left graph) Calcium events rate across intervals (middle graph; n=451 neurons from 3 mice: Wilcoxon signed rank test Baseline-ITI1 *P*<0.002). (Right graph) Decreased rate upon removal of the social stimulus (Wilcoxon signed test T1-ITI1 *P*<0.0001). Data shown as mean±sem: *,*P*<0.05, #,*P*<0.0001
